# The influence of auditory attention on rhythmic speech entrainment

**DOI:** 10.1101/2020.04.30.070318

**Authors:** Rodika Sokoliuk, Giulio Degano, Lucia Melloni, Uta Noppeney, Damian Cruse

## Abstract

Language comprehension relies on integrating words into progressively more complex structures, like phrases and sentences. This hierarchical structure building is reflected in rhythmic neural activity across multiple timescales in E/MEG (Ding et al., 2016, 2017).

How does selective attention across levels of the hierarchy influence the expression of these rhythms?

We investigated these questions in an EEG study of 72 healthy human volunteers listening to streams of monosyllabic isochronous English words that were either unrelated (scrambled condition) or composed of four-word-sequences building meaningful sentences (sentential condition). Importantly, there were no physical cues between four-word-sentences but boundaries were marked by syntactic structure and thematic role assignment. Participants were divided into three attention groups: from passive listening *(passive group)* to attending to individual words *(word group)* or sentences *(sentence group).* The *passive* and *word group* were naïve to the sentential structure of the stimulus material, while the *sentence group* were not.

We found significant entrainment at word- and sentence rate across all three groups, with sentence entrainment linked to left middle temporal gyrus and right superior temporal gyrus. Goal-directed attention to words did not enhance word-rate-entrainment suggesting that word entrainment relies on largely automatic mechanisms. Importantly, goal-directed attention to sentences relative to words significantly increased sentence-rate-entrainment over left inferior frontal gyrus. This attentional modulation of rhythmic EEG activity at the sentential level highlights the role of attention in integrating individual words into complex linguistic structures.

**SIGNIFICANCE STATEMENT:** Neural activity is known to entrain to physical characteristics of auditory stimuli. However, entrainment also occurs with structures lacking physical cues but rather require comprehension of the stimulus’ meaning – for example, entrainment to sentences in speech even without acoustic gaps separating these higher linguistic structures.

We investigated how goal-directed attention to low-level (words) and high-level (sentences) linguistic structures influences entrainment strength. Whilst sentence entrainment occurred independently of selective attention, it increased with goal-directed attention towards sentences. Conversely, no such attentional effect was found for word entrainment.

While goal-directed attention towards sentences strengthens entrainment, it is no prerequisite for it to occur, suggesting that low attentional effort is required for sentence comprehension, potentially reflecting the importance of speech in humans.

## INTRODUCTION

Reading a book on public transport can be challenging, especially if people around us talk about their personal life, or the latest gossip in the neighbourhood, leaving us unwillingly trapped in their conversations. Disconnecting becomes easier if we do not understand the language. In both cases, however, our auditory neurons follow the rhythm of individual syllables (4-8Hz; (Ghitza, 2013)), suggested to reflect entrainment of underlying brain oscillations to acoustic features of speech (Poeppel, 2003; Ghitza, 2011, 2012, 2013; Giraud and Poeppel, 2012) boosting its intelligibility (Luo and Poeppel, 2007; Elliott and Theunissen, 2009; Zoefel and VanRullen, 2015, 2016).

Understanding a language is required for additional, slower rhythms, reflecting the semantic link between syllables. For instance, the mono-syllabic words ‘sharp-knife’ are in themselves informative but also build a meaningful phrase (Chomsky, 1957). In the same way, but on a larger scale, we know when a sentence begins and ends based on the meaning of individual syllables/words and their syntax. Importantly, acoustic cues, like gaps separating phrases or sentences, are not necessary. Slow neural rhythms in speech comprehension have been observed with magnetoencephalography (MEG; (Ding et al., 2016)), surface and intracranial electroencephalography (EEG (Ding et al., 2017), iEEG (Ding et al., 2016)). In a MEG study, participants listened to mono-syllabic isochronous words, played consecutively. These were either unintelligible syllables/unrelated words *(scrambled condition;* e.g. *‘cold-eat-cell-dog…’)* or sequences of four syllables/words building a meaningful sentence *(sentential condition;* e.g. *‘sharp-knife-cuts-meat…*’). Both conditions led to a peak at the syllable/word frequency in the power spectra – i.e. entrainment – but crucially, entrainment at the rate of the phrases *(‘sharp-knife’)* and sentences was observed only in the *sentential condition,* despite there being no acoustic stimulus changes at those rates. Importantly, these peaks were only found when participants understood the language (Ding et al., 2016) and were awake (Makov et al., 2017). Therefore, it has been suggested that these slower brain oscillations might reflect speech comprehension (Ding et al., 2016, 2017; Makov et al., 2017).

However, all of the above studies informed participants about the sentential stimulus structure and involved an active task based on sentences. Would sentence entrainment also be present in naïve participants and without an active task on sentences? What is the influence of goal-directed attention on acoustic/linguistic entrainment?

We investigated these questions in an EEG study on 72 healthy human volunteers, listening to mono-syllabic isochronous English words (cf. Figure 1). Similar to the previous studies, in the *sentential condition,* four consecutive words built a meaningful sentence, whereas in the *scrambled condition*, successive words were unrelated. Participants were either passively listening (*passive group*), attending to individual words (*word group*), or to sentences (*sentence group;* comparable with previous studies). While the *passive* and *word group* were naïve to the sentential stimulus structure, the *sentence group* was instructed about it prior to the experiment. This combination of stimulus and task manipulations allowed us to orthogonally isolate the roles of attention on neural entrainment to low-level (words) and high-level (sentences) features.

**Figure 1:**
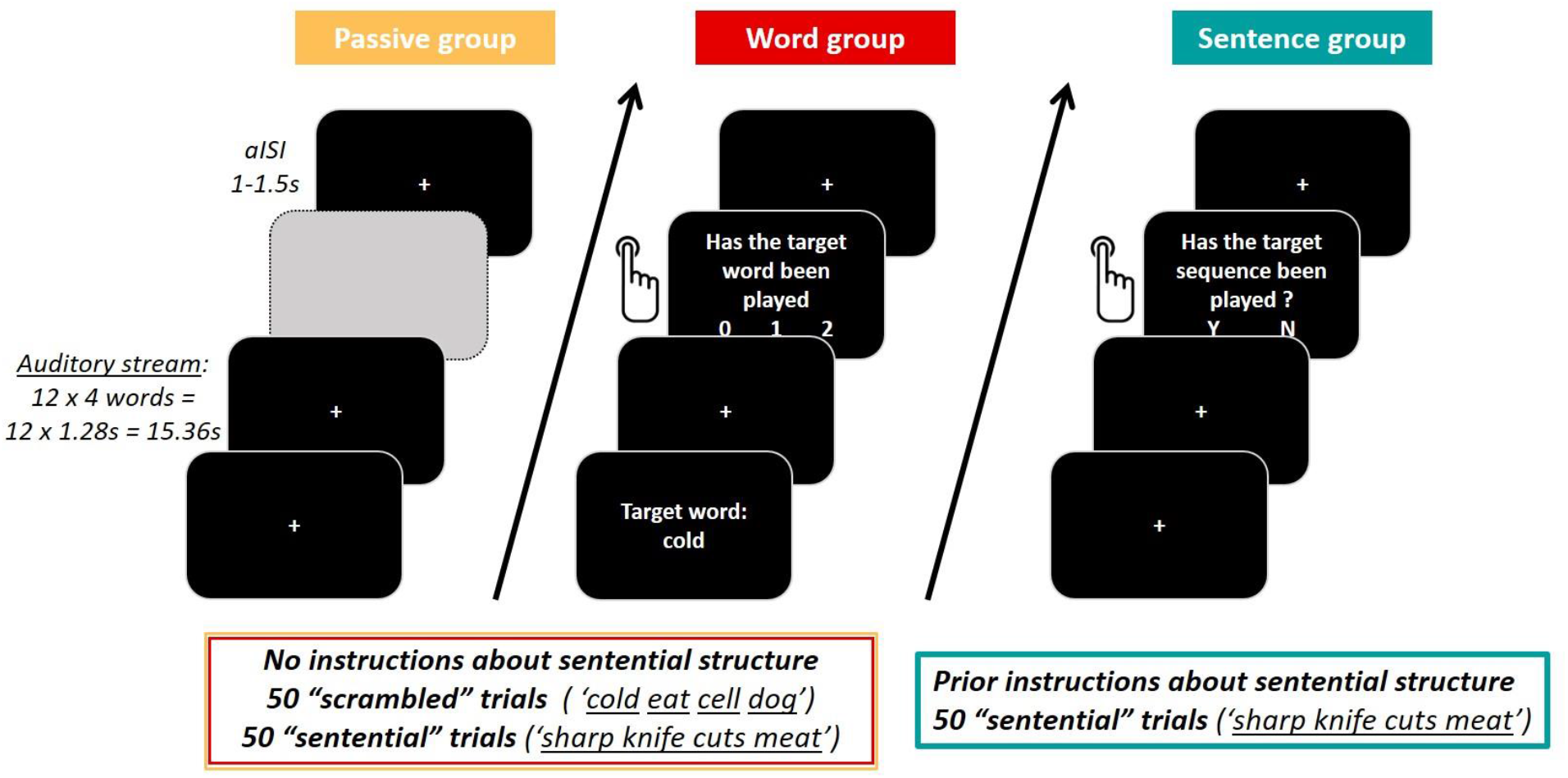
Experimental paradigm. Participants were divided into 3 groups of 24 participants each. For every group, auditory stimuli of each trial were presented as a continuous stream of twelve 4-word-sequences, which were built by concatenating isochronous single-words of 320ms length each. The passive group (orange) was naïve to the sentential structure of the stimulus material and passively listened to the auditory stream. The word group (red) was also naïve to the sentential structure and had an active task based on individual target words. After each trial, participants were asked whether the target word had been presented 0,1, or 2 times. Both the passive and the word group were exposed to stimuli of the scrambled condition (unrelated words) and the sentential condition (four consecutive words built a meaningful sentence). The sentence group (turquoise) was instructed about the sentential structure in the stimulus material and was only listening to the sentential condition. Participants were asked to report via button press after each trial, whether a grammatically incorrect four-word-sequence had been played in the respective trial. For each participant group, individual trials were separated by an asynchronous inter-stimulus-interval (aISI) of 1-1.5s. (Greyed out field only serves visualisation purposes of this figure.)

All groups showed significant entrainment to words, phrases, and sentences. In the *sentence group,* goal-directed attention to sentences led to significantly increased sentence entrainment strength. No such attentional effect was observed at the word rate for the *word group* compared with the other groups, despite their goal-directed attention to individual words.

The sentence entrainment across all groups suggests it neither requires explicit attention to the sentential structure nor prior knowledge about it. Yet, goal-directed attention does strengthen the sentence entrainment.

We therefore replicate previous results (Ding et al., 2016, 2017) and show that sentence-rate entrainment is not simply a result of goal-directed internal attentional sampling (every four words) but also observable without selective attention to sentences. These findings illustrate how we are committed to listening to a language we understand, even if we prefer not to.

## MATERIALS & METHODS

### Participants

We recorded behavioural and EEG data of 72 healthy human volunteers (median age: 22, range: 18-33; 42 females). All participants reported to be monolingual native English speakers, between 18 and 35 years old, right-handed, with no history of epilepsy, and no diagnosis of dyslexia. Participants received either course credits or a monetary compensation for participation. The experimental procedures were approved by the Ethical Review Committee of the University of Birmingham (ERN_15-1367AP3) and conformed with the Declaration of Helsinki. All participants gave written informed consent prior to participation in the study. Data of five participants were excluded from the analysis because of excessive artefacts in the EEG signal and two participants in the sentence group because of poor behavioural performance (<=50%), resulting in total in 20, 23 and 22 participants for the *passive group,* the *word group,* and for the *sentence group,* respectively. Participants received either course credits or monetary compensation for participation.

### Stimuli

We constructed a total of 288 mono-syllabic English words using the male voice of the Apple synthesiser (Macintalk, voice Alex; Apple MacBook Pro Third generation) and these words were segmented using Audacity software version 2.1. Importantly, words were isochronous, of 320ms in length, which resulted in a presentation frequency of 3.125 Hz for the word rate, 1.56Hz for the phrase rate and 0.78Hz for the sentence rate. The words included 144 nouns, 72 adjectives and 72 verbs (full word list is available on OSF under the following link: https://osf.io/8pu4a/?view_only=989f97d6acc44065b052752f00b4ee06). For the *sentential* condition, a total of 72 four-word-sentences were constructed, conforming to the syntactic structure: adjective – noun – verb – noun. Each four-word-sentence was played a minimum of eight and a maximum of nine times per participant throughout the experiment. The order with which they were presented was randomly chosen on a trial-by-trial basis, avoiding occurrence of the same four-word-sentence more than once per trial. For the sentence group only, 10% of the trials contained target sequences, which were grammatically incorrect and either followed the order ‘noun – noun – adjective – verb’ or ‘adjective – verb – noun – noun’.

In the *scrambled* condition, every trial consisted of twelve four-word-sequences. Each contained four randomly chosen individual words of a given word category, i.e., noun, adjective and verb from the total of 288 words. Every word was played a minimum of 15 and a maximum of 50 times. To ensure that no grammatically correct sequences were presented, half of the scrambled sequences followed the order ‘noun – noun – adjective – verb’, the other half followed the order ‘adjective – verb – noun – noun’.

The *sentential* and *scrambled* condition both contained 50 trials with twelve four-word sequences each resulting in a total of 600 scrambled four-word-sequences and 600 meaningful four-word-sentences.

Throughout the experiment, participants were instructed to fixate a white cross at the centre of the screen, to minimise ocular as well as head movements. All stimuli were presented via the MATLAB toolbox Psychtoolbox (Brainard, 1997).

To validate the stimulus material, we performed a control analysis to ensure that the entrainment to higher linguistic structures such as phrases or sentences did not reflect entrainment to the acoustic envelope of the material. Therefore, we ran a bootstrap analysis, creating a random set of 50 trials per participant (random selection with replacement; 50 trials corresponding to the number of trials in our study) of the existing EEG and acoustic data per repetition (1000 repetitions). ITPC measures were then computed on auditory and EEG data per repetition, ITPC values at target frequencies extracted (0.78Hz (sentences), 1.56Hz (phrases) and 3.125Hz (words)) and averaged over repetitions and participants. In a Monte Carlo test, these average ITPC values at target frequencies were then compared to the distribution of ITPC values at “chance frequencies” (i.e. 500 non-harmonic frequencies of the target frequencies) – which were also averaged over repetitions and participants – for acoustic and EEG data. The results of this analysis revealed that only the EEG data showed significant ITPC values at all target frequencies (p<5×10^-4^) whereas acoustic data showed a significant ITPC value only at the word rate (p<5×10^-4^) but not at the phrase- (p=0.502) or sentence rate (p=0.505). We can therefore conclude that there is no information in the acoustic envelope of the stimulus at either of the higher-level linguistic rates.

### Experimental design

The experiment included three groups of subjects.

The *passive group* was naïve to the sentence structure and only passively listened to the stimulus material.

The *word group* was also naive to the sentence structure with their attention being directed to the individual words. After each trial (i.e. twelve four-word-sequences), they judged whether a particular target word which was presented on the screen before each trial, appeared zero, one or two times within the auditory stream of the previous trial. Target words were either adjectives or verbs (50% adjectives and 50% verbs) randomly chosen from the pool of words used in this paradigm. A total of 10% of the trials contained target words.

Both *passive* and *word group* were presented with scrambled and sentential word sequences. The *sentence group* was presented with the sentential condition only where 10% of the sequences were grammatically incorrect sentences. Participants were informed about the sentential stimulus structure prior to the experiment and were asked to perform a task based on these sentences: they had to identify the grammatically incorrect four-word-sequences (e.g. *cold-eat-cell-dog),* by responding after each trial (i.e. twelve four-word-sequences) with ‘yes’ or ‘no’.

For all groups, individual trials were separated by a jittered delay of 1-1.5s (cf. asynchronous inter-stimulus-interval (aISI) in Figure 1).

### Procedure

During the study, participants sat comfortably in a dim room, ~50cm in front of an LCD screen. The experiment was divided into five blocks. Participants self-initiated a block by pressing a button on the keyboard. They were instructed to take breaks in between blocks if needed. Each block included 20 trials for *passive* and *word group* (i.e. sentential and scrambled conditions) and 10 trials for the *sentence group* (only sentential condition), where each trial consisted of 12 four-word-sequences.

All three participant groups were asked to fixate the central fixation cross throughout the experiment, however received different task instructions (cf. above: *Experimental design).*

After the experiment, participants of the *passive* and the *word group* were asked whether they noticed something specific about the stimulus material without informing them about the sentential structure. This way, we assessed information about whether participants noticed the sentential structure of the stimuli even without any prior knowledge about it. All participants were further debriefed about the study.

### Behavioural data analysis

We report median, minimum and maximum performance for the word and sentence tasks in the respective groups. Participants whose average performance accuracy was not better than chance, were excluded from data analysis.

To assess whether the trial type influences performance accuracy for the *word group*, their performance accuracy was split between sentential and scrambled trials and the averages over these conditions were compared in a paired t-test.

### EEG data acquisition

EEG data was recorded at 1000Hz via the software eego64 (ANT Neuro, The Netherlands), using a 124-electrode ANT EEG system (ANT Neuro, The Netherlands) with an extended 10/20 layout. The ground electrode was placed on the left mastoid whereas the reference electrode was located at CPz. All electrodes showed an impedance of <20kΩ before the recording started. Individual electrode locations as well as fiducials (nasion, right and left interauricular points) were recorded prior to the experiment using the software Xensor (ANT Neuro, The Netherlands).

### EEG pre-processing

EEG data pre-processing was performed using custom-written Matlab scripts (all analysis scripts can be found under the OSF repository following this link: https://osf.io/8pu4a/?view_only=989f97d6acc44065b052752f00b4ee06) and functions of the Matlab toolbox FieldTrip (Oostenveld et al., 2011). EEG data was filtered between 0.01 and 170Hz, using a FIR filter at filter order 3. Additionally, a notch filter was applied at 48-52Hz, 98-102Hz, and 148-152Hz using a FIR filter to reduce line noise. Subsequently, the data was epoched into trials starting 1s before stimulus onset and lasting for the whole length of each auditory stream. This way, trials of 16.36s were created. Then, data was visually inspected for artefacts as well as noisy channels, which were removed from the data before an ICA was computed (Bell and Sejnowski, 1995) on the down-sampled data (500Hz), to remove blinks and horizontal eye movements from the data. Finally, noisy channels were interpolated by using data of their neighbours, which were identified via the triangulation method, as implemented in FieldTrip (Oostenveld et al., 2011), before the data was re-referenced to average while reconstructing the reference channel, CPz.

Subsequently, a low-pass filter at 25Hz (butterworth) was applied to the data given the low cutoff of the frequencies of interest (<4Hz). In preparation for the next analysis step, all trials were further cut to discard the first 2.28s (resulting in 11 out of the 12 four-word sequences per trial), which correspond to the 1s pre-stimulus period and the first four-word-sequence, to avoid including the transient EEG response to the onset of the auditory stimulus (cf. (Ding et al., 2017).

### EEG analysis: sensor-level

Inter-trial-phase-coherence (ITPC) was used as a measure to quantify whether the brain signal carried signatures of the rhythmic auditory stimulation. This was achieved by first computing the Discrete Fourier Transform (DFT) of the data, for each trial and electrode separately, to transform the signal into the frequency domain with 0.07Hz resolution (i.e. 1/(15.36s-1.28s)). ITPC was then computed by extracting the absolute values of the mean over the phase angles of each complex-valued Fourier coefficient (cf. (Ding et al., 2017)). For the *passive* and the *word group,* this was done separately for the sentential and the scrambled condition. ITPC values were further averaged over trials within each condition and participant which resulted in 7041 ITPC values for each of the 125 electrodes (i.e. 7041 frequencies x 125 electrodes) per participant and condition.

### Statistical analysis of EEG data

#### Average ITPC at target frequencies

To test whether each participant group showed significant entrainment at the target frequencies, ITPC values were averaged over all electrodes to obtain one average ITPC value per frequency and condition for each participant. Paired t-tests were computed for each participant group separately, comparing the ITPC values at one of the target frequencies (word rate (3.125Hz), phrase rate (1.56Hz) and sentence rate (0.78Hz)) with the ITPC values averaged over ±7 surrounding frequencies, which corresponds to ±0.5Hz (cf. Ding et al, 2017). This way, potential significant peaks (p<0.05) at the target frequencies could be identified. The resulting p-values were further corrected within each group and for each condition for multiple comparisons (number of target frequencies) by applying a False Discovery Rate (FDR) correction (Benjamini and Hochberg, 1995; Yekutieli and Benjamini, 1999).

#### Scalp distribution analyses

In order to estimate the scalp distribution of the effects of interest, ITPC values across all electrodes were compared with the cluster mass method of the Matlab toolbox FieldTrip (Oostenveld et al., 2011). Briefly, this involves that adjacent electrodes were grouped in a cluster if their t-test p-values passed the threshold (detailed below), with the minimum number of electrodes within a cluster set to 4 (adjacent electrodes were identified using the triangulation method). To correct for multiple comparisons, 1000 Monte Carlo permutations of the above method were produced by a randomisation procedure to estimate the probability of the electrode cluster under the null hypothesis (as implemented in the FieldTrip toolbox (Oostenveld et al., 2011)).

##### a) ITPC scalp distribution specific for listening to sentences

To investigate the scalp distribution of ITPC specific for listening to sentences, ITPC scalp distributions at the sentence rate (0.78Hz) were compared between the *sentential* and the *scrambled* condition. To that end, we pooled together the data from the *passive* and the *word group*. The *sentence group* was not included, since participants were not exposed to the scrambled condition. Since we expected stronger ITPC at the sentence rate (0.78Hz) for the sentential compared with the scrambled condition, we applied a one-tailed dependent samples t-test at each electrode and the alpha level and cluster alpha level were set to 0.05.

##### a) ITPC scalp distribution specific for goal-directed attention to words and sentences

To test for an effect of attention to individual words on ITPC strength, ITPC scalp distributions at the word-frequency (i.e. 3.125Hz) of the sentential condition were compared between the *word* and the *passive group* and between the *word* and the *sentence group,* applying two-tailed independent samples t-tests at each electrode. The alpha level and cluster alpha level were set to 0.025, as here we tested for both, positive and negative electrode clusters.

To investigate the effect of attention to sentences on the entrainment strength at the sentence frequency (0.78Hz), ITPC scalp distributions of the *sentence group* were compared to those of the sentential condition pooled over participants of the *passive* and the *word group.*

The alpha level and cluster alpha level were set to 0.025 as here we tested for both positive and negative electrode clusters.

#### ANOVA: Influence of attention on entrainment strength at different target frequencies

In order to analyse a potential interaction between attention condition and entrainment strength, ITPC values at word (3.125Hz) and sentence frequency (0.78Hz) were compared between participants of the *word* and the *sentence group* using a two-way ANOVA, where attentional manipulation (attention to individual words (*word group*) vs. attention to whole sentences (*sentence group*)) and entrainment frequency (word- and sentence rate) were the independent variables and ITPC was the dependent variable. Therefore, first, for every participant of the *word* and *sentence group*, individual peak electrodes were defined as those electrodes that showed the maximum ITPC value at the word (3.125Hz) and sentence (0.78Hz) frequency respectively. Second, these peak ITPC values then served as input for the ANOVA.

#### EEG sensor-level Bayesian t-tests

To assess evidence supporting the Null hypothesis in the word rate ITPC contrast between *word* and *passive group* as well as between *word* and *sentence group,* we computed Bayesian equivalent 2-sample t-tests. We therefore computed a Jeffrey-Zellner-Siow Bayes factor (JZS-BF) at each electrode, as implemented in an open-access script (https://github.com/anne-urai/Tools/tree/master/stats/BayesFactors). JZS-BF>3 reflect substantial evidence in support of the tested hypothesis, while JZS-BF<0.33 reflect substantial evidence in favour of the Null hypothesis.

### EEG/MRI co-registration

We recorded the electrode locations of each participant relative to the surface of the head using the infrared camera system device *Xensor* (ANT Neuro, The Netherlands). Because we did not acquire individual T1-weighted MRI images for all our participants, we used the template files provided by FieldTrip (MRI file, headmodel and grid) and co-registered the standard T1-weighted anatomical scan of the FieldTrip template (1mm voxel resolution) to the digitised electrode locations using Fieldtrip (Oostenveld et al., 2011).

### EEG source estimation

All source analyses were carried out using Dynamic Imaging of Coherent Sources (DICS (Gross et al., 2001)) beamforming.

#### Source estimation specific for listening to sentences

To estimate the sources specific for listening to four-word-sentences, we pooled together data of participants of the *passive* and the *word group* and compared sentence entrainment between the sentential and the scrambled condition.

Therefore, we first computed the cross-spectral density matrix at the sentence frequency (0.78Hz). We therefore used the method ‘mtmfft’ with spectral smoothing of ±0.071Hz and cross-spectral density matrix and power as output, as implemented in FieldTrip. We did that for each trial of the sentential and scrambled condition separately as well as for the combined data (sentential and scrambled trials together). The cross-spectral density matrix of the combined data then served as input to create the common spatial filter for this contrast.

Second, we computed the common spatial filter (regularisation parameter = 5%) which was applied to the cross-spectral-density matrix of the individual conditions. We then contrasted the power of the sentential condition with the power of the scrambled condition in source space and visualised the results on a standard MNI brain using the visualisation software caret (Van Essen and Drury, 1997).

#### Source estimation of attention effect on ITPC

To investigate the effect of attention on ITPC strength at the sentence frequency on the source level, we compared sentence rate entrainment between the *sentence* and the *word group.*

Therefore, we first computed the cross-spectral density matrix at the sentence frequency (0.78Hz) for each trial of the sentential condition of each participant of the *sentence* and the *word group*, using the same parameters as described above.

Second, we computed a spatial filter for the sentence entrainment for every participant (5% regularisation parameter). Since for this contrast, only one experimental condition was investigated per participant, we computed the *neural activity index* (NAI) by normalising the obtained source results of each participant by the estimated noise obtained from the source analysis. This approach has been shown to circumvent the noise bias towards the center of the brain. Subsequently, the NAI source estimates were contrasted between participants of the *sentence* and the *word group* and visualised on a standard MNI brain using the visualisation software caret (Van Essen and Drury, 1997).

## RESULTS

### Behavioural Results

All participants of the *word* and 22 out of 24 participants in the *sentence group* performed the behavioural task above chance *(word group:* chance level: 33.33%; median performance: 87%, range: 72-96%; *sentence group:* chance level: 50%; median performance: 80%, range: 45-95%); two participants had to be excluded from the sentence group based on this criterion (median performance after exclusion of this participant: 81.25%, range: 55-95%). Participants of the *word group* further showed a benefit of the sentential condition on identifying target words compared with the scrambled condition (T(22) = 7.173; p = 3.438 x 10^-7^; median performance_sentential_: 94%, range: 78-100%; median performance_scrambled_: 80%, range: 62-92%). Furthermore, debriefing of participants of the *passive* and the *word group* preceding the experiment revealed that all participants noticed the sentential character of the auditory material of the sentential condition, without any prior instruction.

#### EEG Results – Significant entrainment at all levels for all participant groups

The ITPC values showed significant peaks at all target frequencies for each of the participant groups (Figure 2; *passive group:* orange, *word group:* red, *sentence group:* turquoise) in the sentential condition (subsequent FDR-corrected p-values reported in the order: *passive, word* and *sentence group;* word rate (3.125Hz): T(19) = 10.765, p= 9.5×10^-9^, T(22) = 13.141, p= 4.1×10^-11^, T(21) = 11.413, p= 5.5×10^-10^; phrase rate (1.56Hz): T(19) = 3.061, p= 0.013, T(22) = 5.359, p= 4.4×10^-5^, T(21) = 9.233, p= 1.2×10^-8^; and sentence rate (0.78Hz): T(19) = 2.842, p= 0.016, T(22) = 3.783, p= 0.002, T(21) = 6.082, p= 5×10^-6^) (see Figure 2). The *passive* and the *word group* further showed significant ITPC peaks at the word rate for the scrambled condition (subsequent FDR-corrected p-values reported in the order: *passive* and *word group;* T(19) = 9.700, p= 2.6×10^-8^, T(22) = 11.146, p= 4.9×10^-10^) and no significant peaks for phrase-(T(19) = 2.101, p= 0.060, T(22) = 0.544, p= 0.592) or sentence rate (T(19) = 0.591, p= 0.562, T(22) = −1.362, p= 0.187) (results not shown).

**Figure 2:**
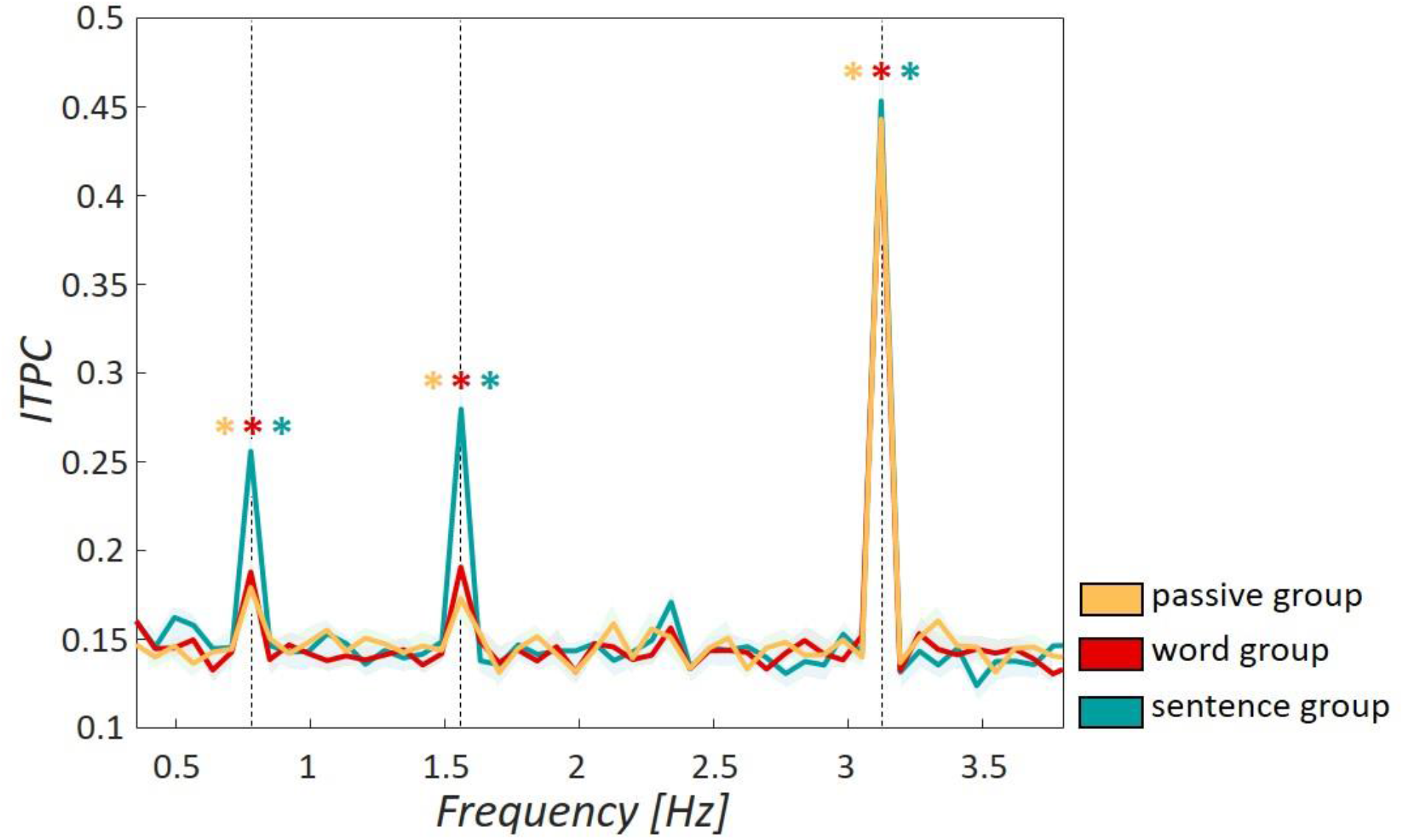
Entrainment to target frequencies over different participant groups. Inter-trial-phase coherence was used as a measure of entrainment strength to the target frequencies at 0.78 Hz (sentences), 1.56Hz (phrases) and 3.125Hz (words). All three participant groups (passive group in orange, word group in red and sentence group in turquoise) showed significant entrainment at all target frequencies (p<0.05). Shaded areas around curves show standard error of the mean; asterisks mark significant entrainment peaks and black dashed vertical lines mark target frequencies. Data of the scrambled condition for participants of the passive and the word group are not shown here.

#### EEG Results – ITPC spatial cluster analysis: Sentential vs. scrambled condition

To test for potential spatial clusters specific to hearing four-word-sentences, we compared ITPC values at the sentence frequency between the sentential and the scrambled condition pooled over participants of the *passive group* and *word group* (Figure 3A). We found a significant positive cluster over left-lateralised parieto-temporal recording sites (Figure 3B) showing significantly stronger entrainment for the sentential relative to the scrambled condition. DICS source estimates of this contrast reveal peaks in the left middle temporal gyrus as well as in the right superior temporal gyrus (Figure 3C).

**Figure 3:**
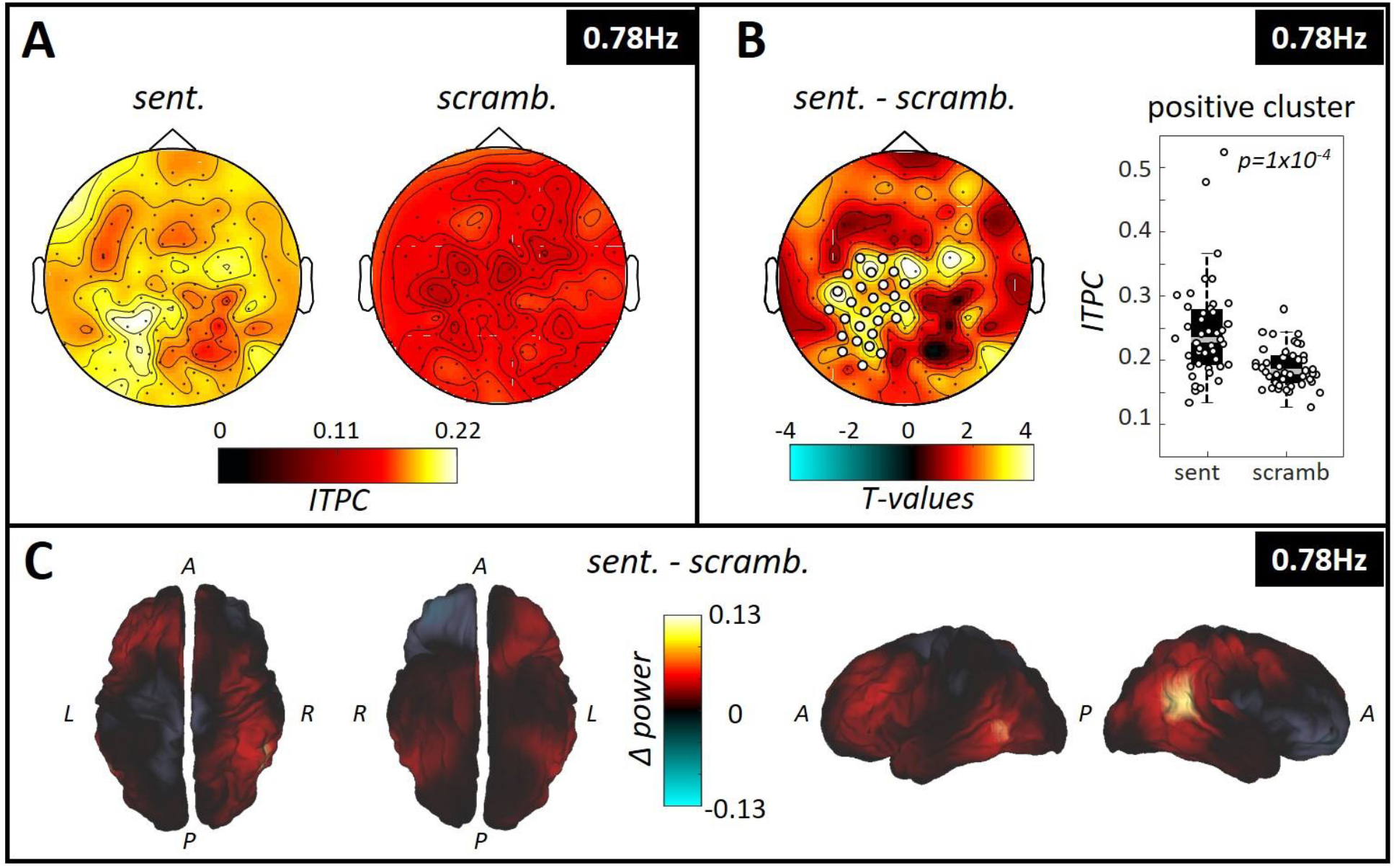
Spatial cluster specific for hearing sentences. (A) Topography plots show colour-coded ITPC values at the sentence frequency (i.e. 0.78Hz) for the sentential (left “sent.”) and the scrambled (right “scramb.”) condition. The data was pooled over all participants of the passive and the word group. (B) When computing the difference between sentential and scrambled condition, a significant positive electrode cluster (p=9.9×10-4) was found, located left-lateralised over parieto-temporal recording sites. The box plot to the right reflects ITPC values over the electrodes of the positive cluster, showing stronger ITPC at the sentence frequency for the sentential compared with the scrambled condition. The central gray line marks the median, the bottom and top edges the 25th and 75th percentiles of the data, respectively. Errorbars extend to the extreme values, excluding outliers and circles represent data of individual participants. (C) Results of the DICS source estimation of the contrast sentential vs. scrambled condition. The source estimation shows colour-coded the difference in power at the sentence frequency between the sentential and the scrambled condition for all subjects of the passive and the word group. The peak areas of the source estimation were identified as left middle temporal gyrus as well as the right superior temporal gyrus. (Letters indicate anatomical landmarks; A=anterior, P=Posterior, L=left, R=right).

#### EEG Results – ITPC spatial cluster analysis: effect of attention on word- and sentence rate entrainment

We investigated whether attending to individual words or sentences modulated the entrainment strength at the word and sentence frequencies selectively in the sentence condition. Indeed, attending to sentences significantly enhanced entrainment at the sentence rate over left-lateralised fronto-temporal recording sites (see Figure 4B). By contrast, goal-directed attention to words did not significantly modulate entrainment at the word rate in the sentential condition (Figure 4A). Likewise, Bayes factors provided robust evidence for the absence of an attentional effect on ITPC values at the word-frequency of the sentential condition (i.e. comparable ITPC values for *passive group, the word group and the sentence group).* Figure 4C shows topographies of these contrasts where electrodes with a BF10 < 0.33 are marked as white-filled circles and reflect substantial evidence for the Null.

**Figure 4:**
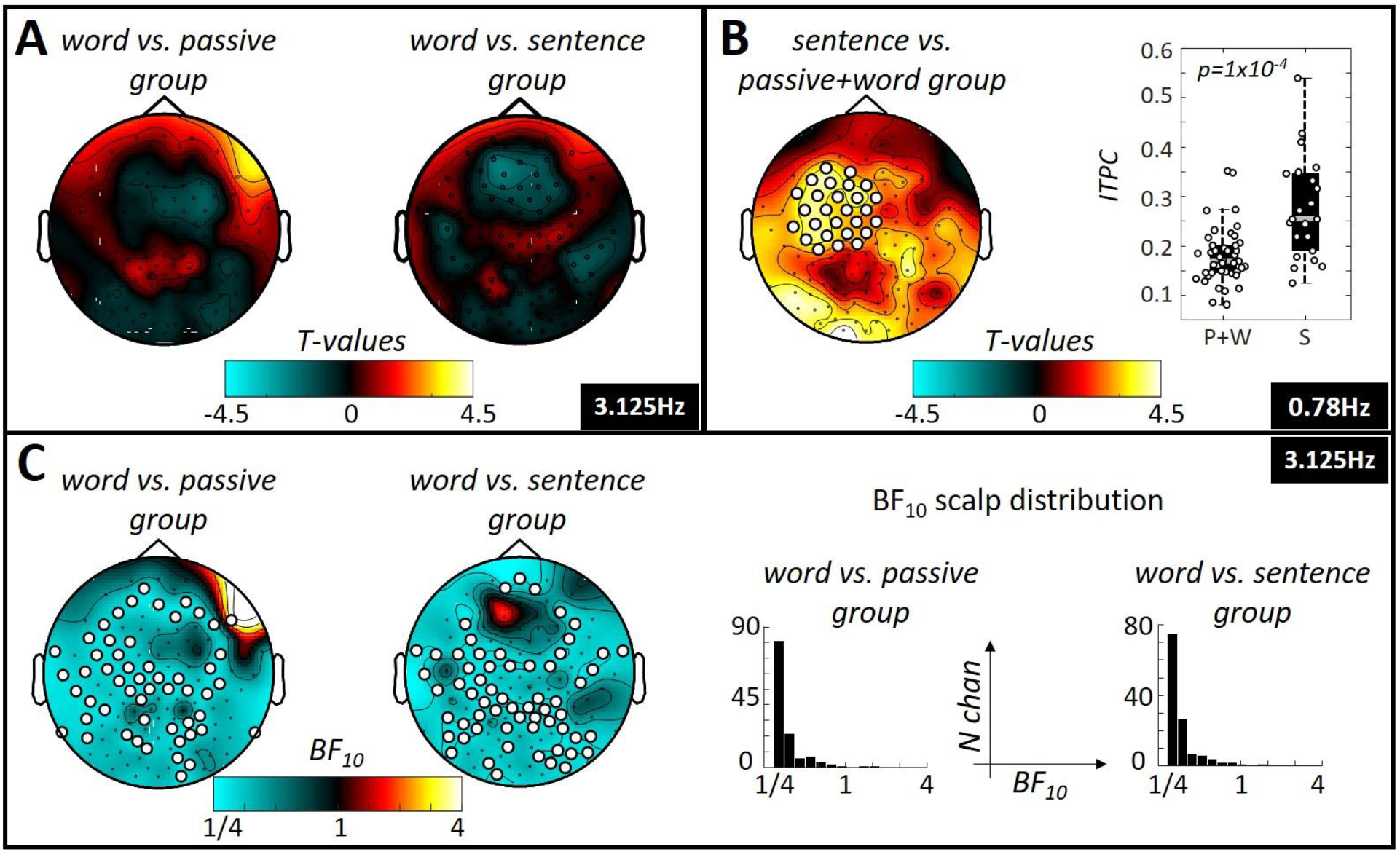
Spatial clusters specific for attentional manipulation. (A) Comparison of the word entrainment strength between participants who paid attention to individual words (word group) with those participants who did not (passive and sentence group). Topography plots show colour-coded T-values of these comparisons. No significant cluster was found. (B) Comparison of the sentence entrainment strength between participants who paid attention to individual sentences (sentence group) with those participants who did not (passive and word group). Topography plot shows colour coded T-values of this comparison. A significant positive cluster was found, located over left-lateralised fronto-temporal recording sites (p=0.002). These clusters are further illustrated in the boxplots on the right side of the panel, showing inter-trial-phase-coherence values for participants of passive + word group and of sentence group for the two clusters. The central gray line marks the median, the bottom and top edges the 25th and 75th percentiles of the data, respectively. Errorbars extend to the extreme values, excluding outliers and circles show data of the individual participants. (C) Results of Bayesian equivalent two-sample t-tests. Topographies show that most electrodes in the contrast word vs. passive group as well as word vs. sentence group show substantial evidence in favour of the Null and thus suggest there is no difference in entrainment strength at the word rate between these groups. Right side of panel (C) shows distributions of BF10 over all electrodes on the scalp for both contrasts.

#### EEG Results – attentional manipulation only shows effect on sentence rate entrainment

A two-way ANOVA with main factors of attention manipulation (i.e. attention to words; attention to sentences) and target frequencies (3.125Hz (word rate) and 0.78Hz (sentence rate)), was computed for ITPC peak electrodes (individually determined for every target frequency and participant). This showed a significant interaction between entrainment frequency and attention condition (F(1,89) = 6.66; p=0.012). Post-hoc t-tests revealed evidence that only entrainment at the sentence rate was modulated by the attentional manipulation, showing significantly stronger inter-trial-phase-coherence values at the sentence frequency for the *sentence group* compared with the *word group* (T(43) = 2.784; p=0.008). ITPC values at the word frequency between the *word group* and the *sentence group* did not significantly differ (T(43) = 0.880; p=0.384; Figure 5 A). Figure 5B shows source estimates of this contrast with a peak in power difference identified over the left inferior frontal gyrus.

**Figure 5:**
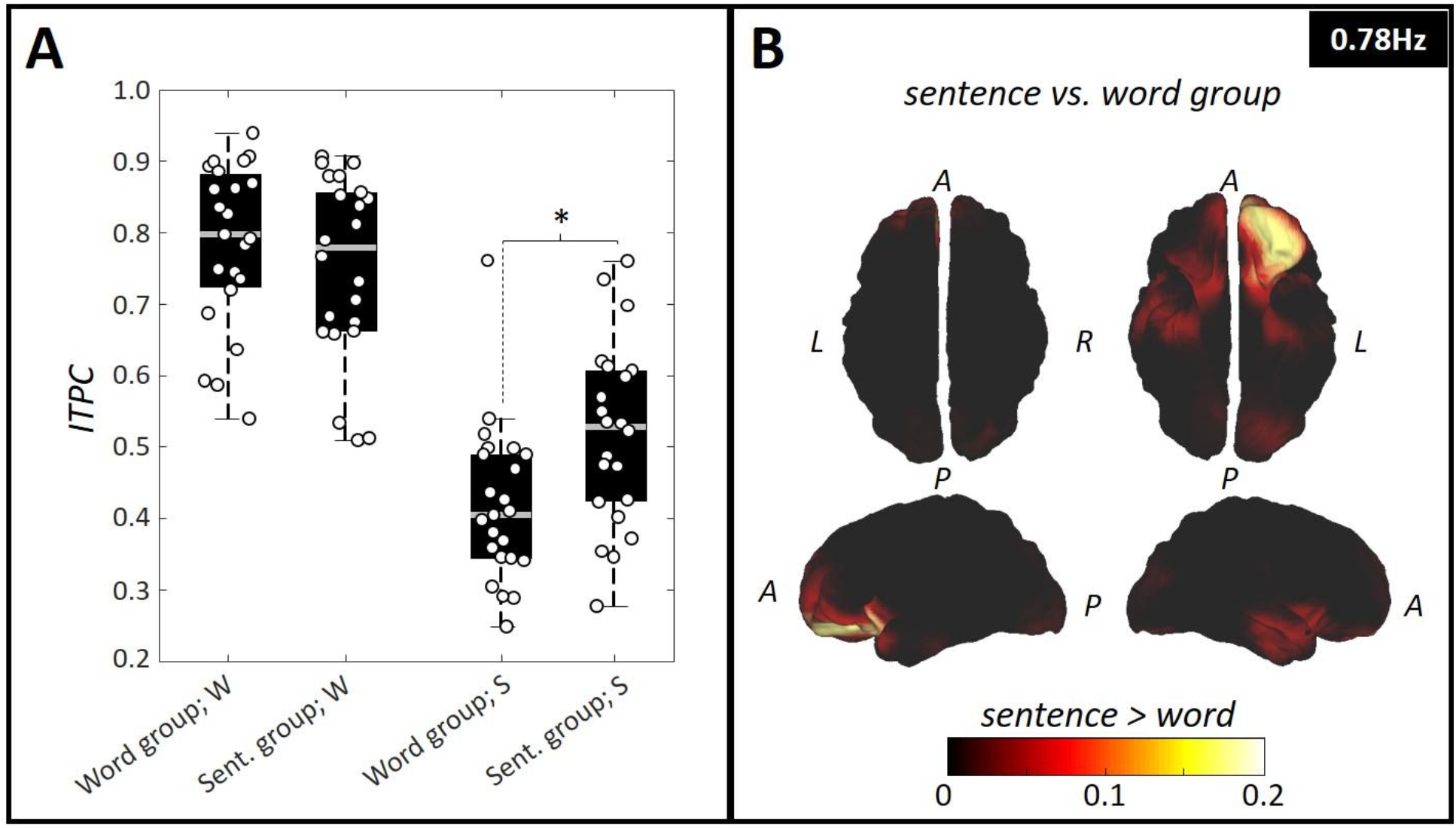
ITPC values at sentence frequency are modulated by attention. (A) Box plots representing ITPC values at word-(“W”) and sentence (“S”) frequency for participants of the word group and the sentence group over the individual peak ITPC electrodes. We observed a significant interaction between entrainment frequency (words (3.125Hz) and sentences (0.78Hz)) and attention condition (attention to words (word group), attention to sentences (sentence group)) (F(1,91) = 5.92; p=0.017). Post-hoc t-tests revealed a significant effect of attention on entrainment strength for the sentence rate entrainment only, showing significantly stronger ITPC values for the sentence group compared with the word group (see, right part of panel (A)). Entrainment strength at the word rate however was comparable between the participant groups (left part of panel (A)). The central gray lines mark the median, the bottom and top edges the 25th and 75th percentiles of the data, respectively. Errorbars extend to the extreme values, excluding outliers and circles show data of the individual participants. (B) Source estimation of difference in power at sentence frequency between sentence and word group. The peak region was identified as left inferior frontal gyrus. (Letters indicate anatomical landmarks; A=anterior, P=Posterior, L=left, R=right).

## DISCUSSION

Previous studies have shown cortical entrainment to higher linguistic structures of speech such as phrases and sentences (Ding et al., 2016, 2017). While low-level entrainment to syllables/words was suggested to reflect the steady state response to the rhythmic acoustic envelope, entrainment to higher-level linguistic structures was interpreted as reflecting conscious speech comprehension (Makov et al., 2017).

In these previous studies, participants were instructed about the sentential stimulus structure and performed active tasks that required sentence comprehension. Therefore, we suggested an alternative hypothesis that phrase- and sentence entrainment could reflect internal attentional sampling of the stimulus material in service of task demands (i.e. ‘Do these two/four words build a meaningful phrase/sentence?’). The findings of our EEG study, however, do not support this hypothesis, as naïve participants who did not perform an active task based on sentences *(passive* and *word group)* still showed significant phrase and sentence entrainment. This higher-level entrainment despite the absence of goal-directed attention could result from implicit attention drawn to the meaningful sentences. Nevertheless, sentence entrainment was significantly enhanced by a task that explicitly required sentence comprehension (i.e. the *sentence group*). This result is therefore consistent with the role of slower, cross-word neural oscillations in speech comprehension.

At the source level, the regions specific for sentence comprehension (Figure 3) were identified as left middle temporal gyrus (MTG) and right superior temporal gyrus (STG). Although the lack of individual head models could have decreased source estimate precision, our observed cortical sources have previously been linked to sentence comprehension. For example, the left MTG is more strongly activated when listening to semantically congruent compared with semantically random sentences (Humphries et al., 2007). Furthermore, the right STG has been shown to be activated during semantic processes at the sentence level (Kuperberg et al., 2000). These source estimates are therefore consistent with a functional role of cross-word entrainment in comprehending the sentence structure of the stimuli.

Our source estimates of the attentional modulation of sentence entrainment revealed the left IFG as peak region, which has previously been shown to be involved in language comprehension tasks (e.g. (Obleser and Kotz, 2010; Giustolisi et al., 2018; Kroczek et al., 2019)). The increased activation of the left IFG in our study in response to increased attention towards sentences could reflect comprehension processes in service of task goals, involving regions higher up the language processing hierarchy (left IFG), rather than comprehension itself, which our data indicate is supported by lower-level regions such as left MTG in the comprehension contrast above. While the left IFG has shown stronger activation specific to sentence entrainment previously (Ding et al., 2016), it has also been shown to be more strongly activated upon listening to complex compared with simple sentences (Caplan et al., 2000; Walenski et al., 2019). Regarding our findings, even though the sentences were the same between groups and did not differ in complexity, the *sentence group* focused more on the syntactic structure to identify grammatically incorrect sequences.

Furthermore, our source estimates and scalp distributions showed that increased attention to sentences does not lead to stronger activation of the regions specific for sentence comprehension themselves, but instead, to recruitment of higher-level cortex (cf. Figure 3 and 5). However, previous studies using iEEG found higher-level cortex (left IFG) to be entrained during sentence comprehension (Ding et al., 2016). While this discrepancy with our results could be caused by inaccurate source localisation in our study, the scalp distributions of our two effects are markedly different (cf. Figure 3 and 5), indicating that not entirely overlapping regions of cortex are implicated in the effects and, therefore, that they reflect dissociable cognitive processes. Indeed, the different task demands in our study likely explain this discrepancy; while participants in the previous studies were instructed about the sentential stimulus structure and performed an active task based on sentences, the participants in our comprehension contrast *(sentences* vs. *scrambled)* were naïve to the stimulus material and either passively listening *(passive group)* or performing a task based on individual words *(word group).* Only when comparing sentence rate entrainment between participants completing a specific sentence task, and those not, did we find increased activation over the left IFG. Together, these results indicate that through the different foci of attention (words/sentences) in our paradigm, we disentangle those regions specifically entrained during sentence comprehension (left MTG/right STG) and those involved in comprehension for task goals.

Interestingly, goal-directed attention to individual words did not increase word rate ITPC in the *word* compared with *passive* and *sentence group*. This is surprising as single words follow a strict rhythm, provoking a steady state auditory response (Regan, 1989), which, according to existing literature, should be stronger with increased attention towards this acoustic feature (Tiitinen et al., 1993; Bidet-Caulet et al., 2007; Müller, 2009; Kim et al., 2011; Bharadwaj et al., 2014). Potentially, word rate ITPC here does not reflect processing of the word’s meaning, as is required by the task, but rather the participants’ non-semantic expectation to hear individual words at a specific rhythm – an expectation that would be the same for all conditions and groups. Furthermore, as words were monosyllabic, it is not possible to separate entrainment to words from entrainment to acoustic envelope. Therefore, we conclude that word-rate entrainment here likely reflects an acoustic process and that an alternative attention manipulation in future studies focused on the acoustic envelope may allow others to observe attentional modulation of this rhythm.

Expectation could also explain the increased entrainment in the *sentence group,* as all sentences follow the same pattern, although, unlike the word task, this requires semantic processing to form the higher-level unit. Indeed, internal temporal tracking of sentence duration would help participants to identify target sequences. Previously, this temporal tracking has been shown to be reflected in the EEG by a steady power decrease, along the time span of a sentence, before increasing again with the beginning of a new sentence (Ding et al., 2016). Nevertheless, the resistance of the word entrainment to word-meaning attentional modulation is consistent with previous evidence that word entrainment strength during sleep was comparable to that of awake and attentive participants (Makov et al., 2017). However, as we observed attentional modulation of entrainment to higher linguistic structures, it is possible that – just as sleep does – a stronger attentional manipulation could abolish higher-level entrainment.

An interesting phenomenon we observed is the ITPC magnitude decrease with increasing linguistic level (words>sentences) (cf. Figure 2). One possible cause could be the frequency with which the different linguistic structures occur in the material. Indeed, words are presented four times as often as sentences. These repetitions could reduce noise in the EEG signal, allowing a more accurate measure of ITPC (Moratti et al., 2007; Vinck et al., 2010; VanRullen, 2016). However, when comparing entrainment strength between full sample datasets at the sentence rate with datasets only containing 25% of the trials at the word rate (i.e. corresponding to words (100%) vs. sentences (25%) representation in stimulus material), sentence was still significantly weaker than word entrainment (T(65)=-17.913; p=1×10^-16^). Therefore, repetition of the linguistic structures (words, sentences) in the stimulus material cannot explain the different entrainment strengths at word- and sentence frequency.

Alternatively, the effect could be linked to cognitive effort. Although we showed that sentence entrainment does not require goal-directed attention to sentences, it clearly requires participants to be awake (Makov et al., 2017), whereas word-rate entrainment was preserved during sleep. We therefore believe the latter demands less cognitive effort than comprehension of sentences, which could explain the increased ITPC at the word frequency. However, future research needs to investigate this question further to make clear assumptions about the cause of the difference in entrainment strength.

In the *word group*, behavioural accuracy to detect target words showed a significant difference between sentential and scrambled condition; showing higher accuracy when stimuli formed four-word-sentences. This is consistent with the role of expectation in language comprehension (Kuperberg and Jaeger, 2016), as hearing connected words that build meaningful entities would allow the listener to predict an upcoming target word based on its context, and so be more likely to be detected, which is not possible in the scrambled condition. Alternatively, chunking the individual words into sentences could improve working memory performance, as whole sentences can be easier remembered than individual words (Gilbert et al., 2015), which may then facilitate reporting target words later.

In conclusion, here we replicated (Ding et al., 2016, 2017) and extended previous findings by characterising the role of auditory attention on cortical tracking of speech stimuli. We observed that entrainment to higher linguistic structures does not depend on and therefore cannot be solely explained by goal-directed attention to sentences but rather reflects a more automatic process. However, goal-directed attention to sentences did significantly increase sentence entrainment in higher-level cortex, consistent with a neural enhancement in service of task demands. The low attentional effort required for sentence entrainment (and comprehension) in this paradigm potentially reflects the importance of speech in humans.

## Acknowledgements

This research was funded by an MRC New Investigator Research Grant to DC (MR/P013228/1).

